# The SARS-CoV-2 Omicron (B.1.1.529) variant exhibits altered pathogenicity, transmissibility, and fitness in the golden Syrian hamster model

**DOI:** 10.1101/2022.01.12.476031

**Authors:** Shuofeng Yuan, Zi-Wei Ye, Ronghui Liang, Kaiming Tang, Anna Jinxia Zhang, Gang Lu, Chon Phin Ong, Vincent Kwok-Man Poon, Chris Chung-Sing Chan, Bobo Wing-Yee Mok, Zhenzhi Qin, Yubin Xie, Haoran Sun, Jessica Oi-Ling Tsang, Terrence Tsz-Tai Yuen, Kenn Ka-Heng Chik, Chris Chun-Yiu Chan, Jian-Piao Cai, Cuiting Luo, Lu Lu, Cyril Chik-Yan Yip, Hin Chu, Kelvin Kai-Wang To, Honglin Chen, Dong-Yan Jin, Kwok-Yung Yuen, Jasper Fuk-Woo Chan

## Abstract

The newly emerging SARS-CoV-2 Omicron (B.1.1.529) variant first identified in South Africa in November 2021 is characterized by an unusual number of amino acid mutations in its spike that renders existing vaccines and therapeutic monoclonal antibodies dramatically less effective. The *in vivo* pathogenicity, transmissibility, and fitness of this new Variant of Concerns are unknown. We investigated these virological attributes of the Omicron variant in comparison with those of the currently dominant Delta (B.1.617.2) variant in the golden Syrian hamster COVID-19 model. Omicron-infected hamsters developed significantly less body weight losses, clinical scores, respiratory tract viral burdens, cytokine/chemokine dysregulation, and tissue damages than Delta-infected hamsters. The Omicron and Delta variant were both highly transmissible (100% vs 100%) via contact transmission. Importantly, the Omicron variant consistently demonstrated about 10-20% higher transmissibility than the already-highly transmissible Delta variant in repeated non-contact transmission studies (overall: 30/36 vs 24/36, 83.3% vs 66.7%). The Delta variant displayed higher fitness advantage than the Omicron variant without selection pressure in both *in vitro* and *in vivo* competition models. However, this scenario drastically changed once immune selection pressure with neutralizing antibodies active against the Delta variant but poorly active against the Omicron variant were introduced, with the Omicron variant significantly outcompeting the Delta variant. Taken together, our findings demonstrated that while the Omicron variant is less pathogenic than the Delta variant, it is highly transmissible and can outcompete the Delta variant under immune selection pressure. Next-generation vaccines and antivirals effective against this new VOC are urgently needed.

**One Sentence Summary:** The novel SARS-CoV-2 Omicron variant, though less pathogenic, is highly transmissible and outcompetes the Delta variant under immune selection pressure in the golden Syrian hamster COVID-19 model.

## INTRODUCTION

Severe acute respiratory syndrome coronavirus 2 (SARS-CoV-2) was first identified in late 2019 and quickly developed into the most important global health challenge in recent decades (*1–3*). Despite rapid generation of vaccines and antivirals, and global implementation of various non-pharmaceutical public health measures, the Coronavirus Disease 2019 (COVID-19) pandemic continues to rage on nearly two years with the emergence of more Variants of Concern (VOCs) that exhibit immune evasion and/or enhanced transmissibility (*4*). The Alpha (B.1.1.7) variant emerged in mid-2020 and quickly outcompeted the Beta (B.1.351) variant (*5,6*). The currently dominant Delta (B.1.617.2) variant with enhanced transmissibility and moderate level of antibody resistance then replaced the Alpha (B.1.1.7) variant since mid-2021.

The recently emerging Omicron (B.1.1.529) variant, first identified in South Africa in November 2021, has now affected more than 100 other countries/regions (*7,8*). This new VOC has an alarmingly high number of mutations (>30) at the spike, which significantly reduce the neutralizing activity of vaccine-induced serum antibodies, as well as therapeutic monoclonal antibodies (*9–15*). Preliminary analyses of the severity of infections caused by the Omicron variant compared to previous variants as determined by hospitalization rates have been inconclusive, with some showing reduced hospitalization rate and others showing a lack of significant difference (*16*). What is more apparent from early epidemiological data is that the Omicron variant is spreading rapidly even in populations with high two-dose COVID-19 vaccination uptake rates (*17,18*). However, whether this is due to the intrinsic transmissibility of the Omicron variant or other extrinsic environmental and social factors is unknown. At present, the *in vivo* pathogenicity, transmissibility, and fitness of this variant are poorly understood. In this study, we investigated these virological attributes of the Omicron variant by comparing them with those of the presently dominant Delta variant in the established golden Syrian hamster model for COVID-19, which closely simulates non-lethal human disease (*19*).

## RESULTS

### The Omicron variant is less pathogenic than the Delta variant in vivo

We first compared the clinical signs, viral burden, and cytokine/chemokine profiles of the Omicron and Delta variants in the golden Syrian hamster model (**Fig. 1A**). The body weight losses (<5%) (P<0.01 to P<0.0001) (**Fig. 1B**) and clinical scores (P<0.05 to P<0.01) (**Fig. 1C**) of the Omicron-infected hamsters were limited, and consistently and significantly milder than those of the Delta-infected hamsters from 2 days post-infection (dpi) to 7 dpi. The Omicron-infected hamsters started to regain their body weights at about 4-5 dpi, which was earlier than the Delta-infected hamsters (**Fig. 1B**). Early after infection (2 dpi), the viral loads and infectious virus titers of the two variants in the nasal turbinate and trachea were similar, but the lung viral loads (P<0.05) and virus titers were about 2 log_10_ units (P<0.0001) lower in the Omicron-infected than Delta-infected hamsters (**Fig. 1D**). During the acute (4 dpi) (**Fig. 1E**) and regenerative (7 dpi) (**Fig. 1F**) phases of infection, the viral burden of the Omicron variant became consistently lower than that of the Delta variant throughout the upper and lower respiratory tract. At 7 dpi, the viral titers in the trachea and lung were already below the detection limit (<100 PFU/mg) in the Omicron-infected hamsters (**Fig. 1F**). Virus shedding of the Omicron variant in the oral swabs (**Fig. S1A**) and feces (**Fig. S1B**) became undetectable by 12 dpi and 14 dpi, which were both earlier than the Delta variant. Corroborative to the viral burden findings, the Omicron-infected hamsters had generally lower tissue cytokines/chemokines gene expression levels between 2 dpi and 7 dpi (**Fig. 1G**). At 7 dpi, the dysregulated inflammatory cytokine/chemokine response has almost completely normalized. The antibody response against variant-specific spike receptor-binding domain (RBD) of the Omicron-infected hamsters was also significantly lower than that of the Delta-infected hamsters (**Fig. S2**).

**Fig. 1.**
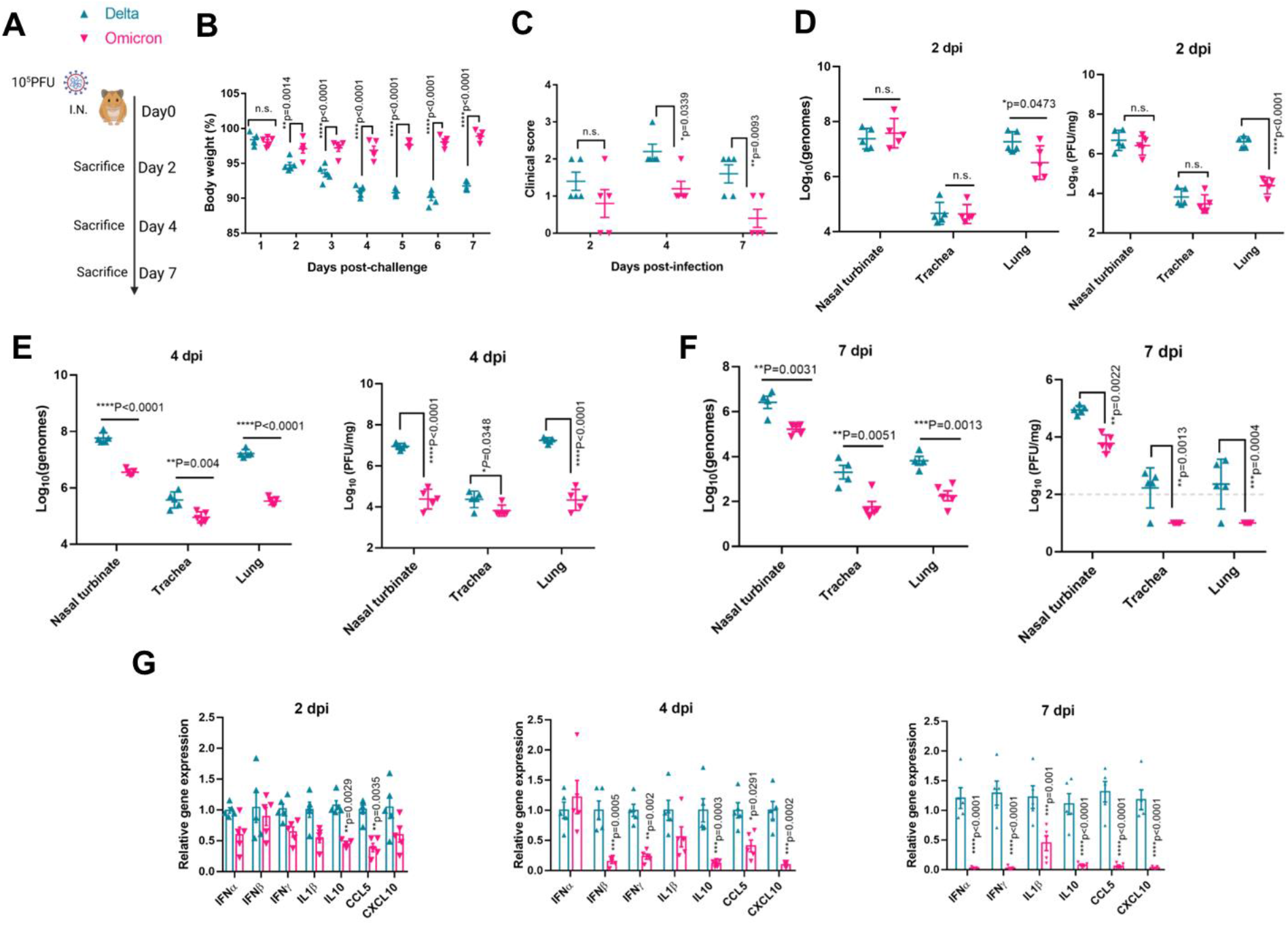
Pathogenicity of the Omicron and Delta variants in the golden Syrian hamster model. **(A)** Scheme of the pathogenicity study comparing infections caused by the Omicron and Delta variants in golden Syrian hamsters. At 0 day post-infection (dpi), each hamster was intranasally inoculated with 100μL of DMEM containing 10^5^ PFU of SARS-CoV-2 (n=15 for each variant). The hamsters were sacrificed at 2dpi, 4dpi, and 7dpi (n=5 per variant per time-point) for viral load quantitation by qRT-PCR, virus titer quantitation by plaque assay, and histopathological studies. **(B)** Body weight changes and **(C)** clinical scores of the hamsters after infection with either variant. A score of 1 was given to each of the following clinical signs: lethargy, ruffled fur, hunchback posture, and rapid breathing. Respiratory tract tissue viral loads (left) and infectious virus titers (right) at **(D)** 2dpi, **(E)** 4dpi, and **(F)** 7dpi (the dotted line represents the limit of detection of the plaque assay at 100 PFU/mg). **(G)** Lung cytokine/chemokine profiles at 2dpi, 4dpi, and 7dpi. Data are mean ± standard deviations. n = 5 biological replicates per variant time-point. \*\*\*\**P*<0.0001, \*\*\**P*<0.001, \*\**P*<0.01, and \**P*<0.05 by two-way ANOVA.

In the lung sections collected at 2 dpi, the Omicron-infected hamsters showed alveolar wall congestion, while the Delta-infected hamsters demonstrated more severe and diffuse alveolar wall infiltration and congestion (**Fig. S3**). At 4 dpi, both groups of hamsters showed bronchiolar epithelial destruction, and peribroncheolar and perivascular inflammatory infiltration, but the alveolitis in the Delta-infected hamsters was more much more diffuse than the Omicron-infected hamsters. At 7 dpi, the lung sections of the Omicron-infected hamsters appeared mostly normal, while those of the Delta-infected hamsters still showed blood vessel congestion and alveolar wall inflammatory infiltration, indicating that the Omicron-infected hamsters had earlier resolution of tissue damage. Viral nucleocapsid protein expression was significantly more abundantly seen in the lung sections of the Delta-infected than the Omicron-infected hamsters throughout 2 dpi to 7 dpi (**Fig. S4**). Similarly, the Omicron-infected hamsters showed less severe histopathological changes (**Fig. S5**) and less abundant viral nucleocapsid protein expression (**Fig. S6**) in their nasal turbinate than the Delta-infected hamsters from 4 dpi to 7 dpi.

### The Omicron variant is highly transmissible through both contact and non-contact routes

The other key question was the comparative transmissibility of the Omicron and Delta variants *in vivo*. To this end, we first co-housed 6 index SARS-CoV-2-challenged hamsters (n=3 each for the Omicron and Delta variants) with 6 naïve hamsters for 4 hours in a 1:1 ratio (**Fig. 2A**). The experiment was repeated twice. All the index hamsters had similar viral loads in the nasal turbinate at sacrifice at 2 dpi, indicating that they were successfully infected (**Fig. 2B**). All 12 naïve hamsters were found to be infected at sacrifice at 2 days after exposure (**Fig. 2C**), indicating that both variants are highly transmissible through close contact. Next, we randomly grouped 42 hamsters into 6 groups of index and naïve hamsters (1:6 ratio) in our established non-contact transmission system, and repeated the experiment twice (total n=84) (**Fig. 2D**) (*20*). The hamsters were sacrificed at 2 dpi (index) or 2 days post-exposure to index (naïve). All the index hamsters were successfully infected with similarly high nasal turbinate virus titers (**Fig. 2E**). A total of 30/36 (83.3%) and 24/36 (66.7%) of naïve animals became infected by the Omicron and Delta variant, respectively, through non-contact transmission (**Fig. 2F**). Notably, although not yet reaching statistical significance with the sample size of 36 naïve hamsters per group (P=0.173), the transmission rate of Omicron variant was consistently about 10-20% higher than that of the Delta variant in both rounds of non-contact transmission experiments. This consistently higher transmission rate would likely be significant in real life scenario with large susceptible human populations.

**Fig. 2.**
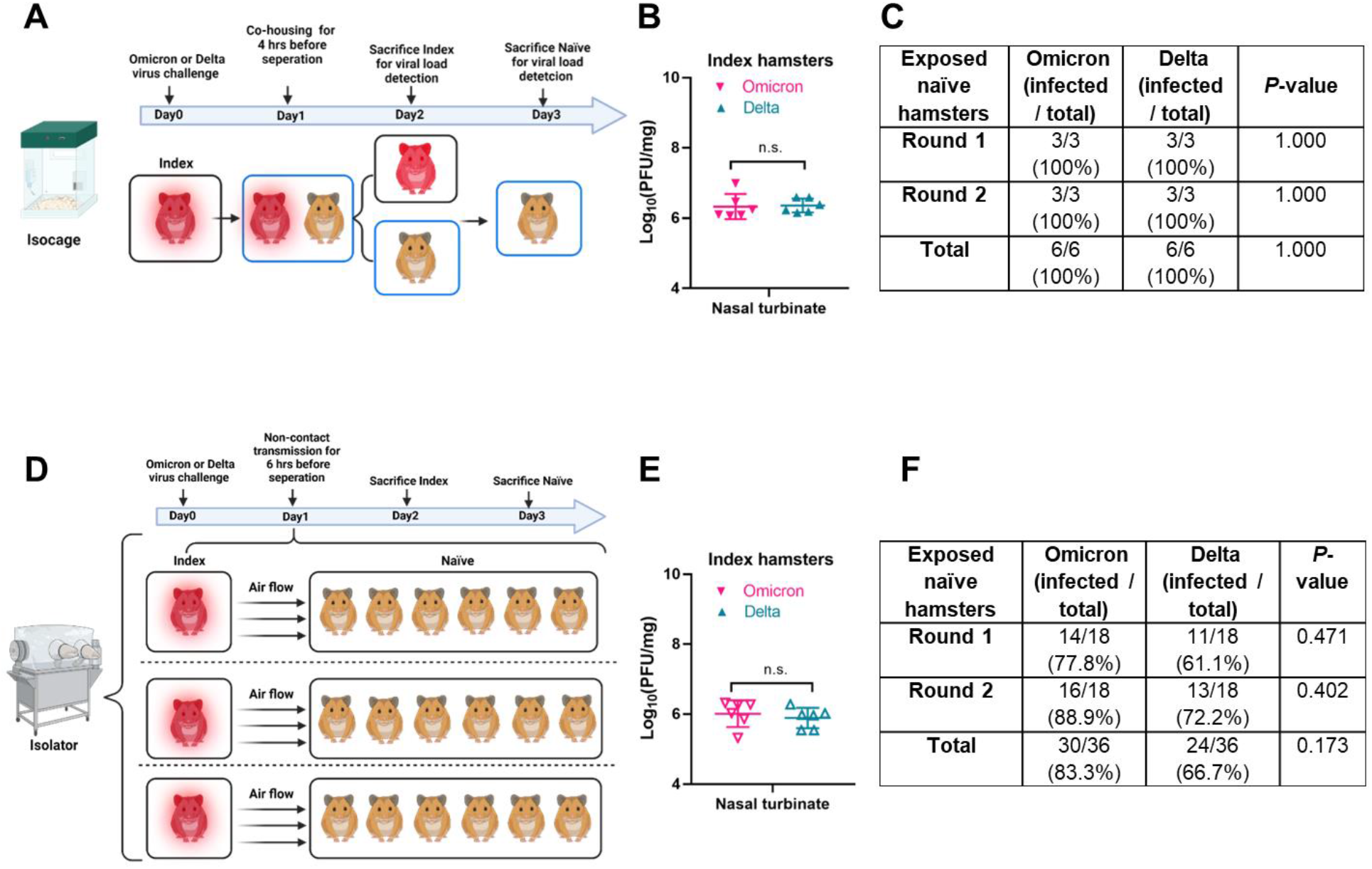
Contact and non-contact transmissions of the Omicron and Delta variants among golden Syrian hamsters. **(A)** Scheme of the contact transmission study. **(B)** The nasal turbinate infectious virus titres in the virus-challenged index hamsters at 2 dpi were determined by plaque assay to ensure successful infection of both groups of hamsters. n.s. indicates non-significant (Student’s t-test). **(C)** Positive rates of infection among the exposed naïve hamsters after exposure to either the Omicron or Delta variant. *P*-values were determined by Chi-square test. **(D)** Scheme of the non-contact transmission study. **(E)** The nasal turbinate infectious virus titres in the virus-challenged index hamsters at 2 dpi were determined by plaque assay to ensure successful infection of both groups of hamsters. n.s. indicates non-significant (Student’s t-test). **(F)** Positive rate of infection after exposure to Omicron or Delta virus. *P*-values were determined by Chi-square test.

### The Omicron variant outcompetes the Delta variant under immune selection pressure

To investigate whether this highly transmissible Omicron variant is likely to take over as the dominant circulating SARS-CoV-2 variant, we performed a competition assay to compare its fitness with that of the presently circulating Delta variant. We co-infected human lung-derived Calu-3 cells with the Omicron and Delta variants. After 24 hours, we collected both culture supernatant and cell lysate, and quantified the relative amounts of RNA of the Omciron and Delta variants by quantitative reverse transcription-polymerase chain reaction (qRT-PCR) and Sanger sequencing. Both methods were validated to reliably quantify the relative amounts of the Omicron and Delta variants in mixed specimens (**Fig. S7**). Consistent with our recent preliminary findings at an early time-point, the Delta variant consistently exhibits significant fitness advantage over the Omicron variant for up to 72hpi *in vitro* (**Fig. 3A and 3B**) (*21*). However, this scenario drastically changed when selection pressure by vaccinated sera containing antibodies with reduced anti-Omicron but preserved anti-Delta neutralizing activity was present (**Fig. 3C**), with the Omicron variant significantly (P<0.0001) outcompeting the Delta variant (**Fig. 3D**). Then, we validated our *in vitro* findings with *in vivo* competition models in hamsters. We included both non-vaccinated (**Fig. 3E**) and vaccinated (**Fig. 3F**) index hamsters, and intranasally challenged them with the two variants (1:1 ratio). The vaccinated index hamsters were at 100 days post-vaccination with an inactivated SARS-CoV-2 vaccine and showed waning serum antibody response in comparison with the peak activity at 28 days post-vaccination (**Fig. 3G**). Their serum neutralizing antibody activity against the Omicron variant was markedly lower than the Delta variant (**Fig. 3H**). In the non-vaccinated index hamsters, the Delta variant significantly outcompeted the Omicron variant (**Fig. 3I**). In stark contrast, the Omicron variant exhibited marked fitness advantage over the Delta variant in the vaccinated index hamsters (**Fig. 3I**). Regarding the naïve hamsters, the Delta variant similarly outcompeted the Omicron variant in those that were exposed to non-vaccinated index hamsters (**Fig. 3J**). Whereas among the naïve hamsters exposed to vaccinated index hamsters, the Omicron variant significantly outcompeted the Delta variant (**Fig. 3J**). Overall, our findings demonstrated that the Delta variant exhibits fitness advantage over the Omicron variant in the absence of selection pressure. Under immune selection pressure, the Omicron variant significantly outcompetes the Delta variant to become the dominant variant causing infection.

**Fig. 3.**
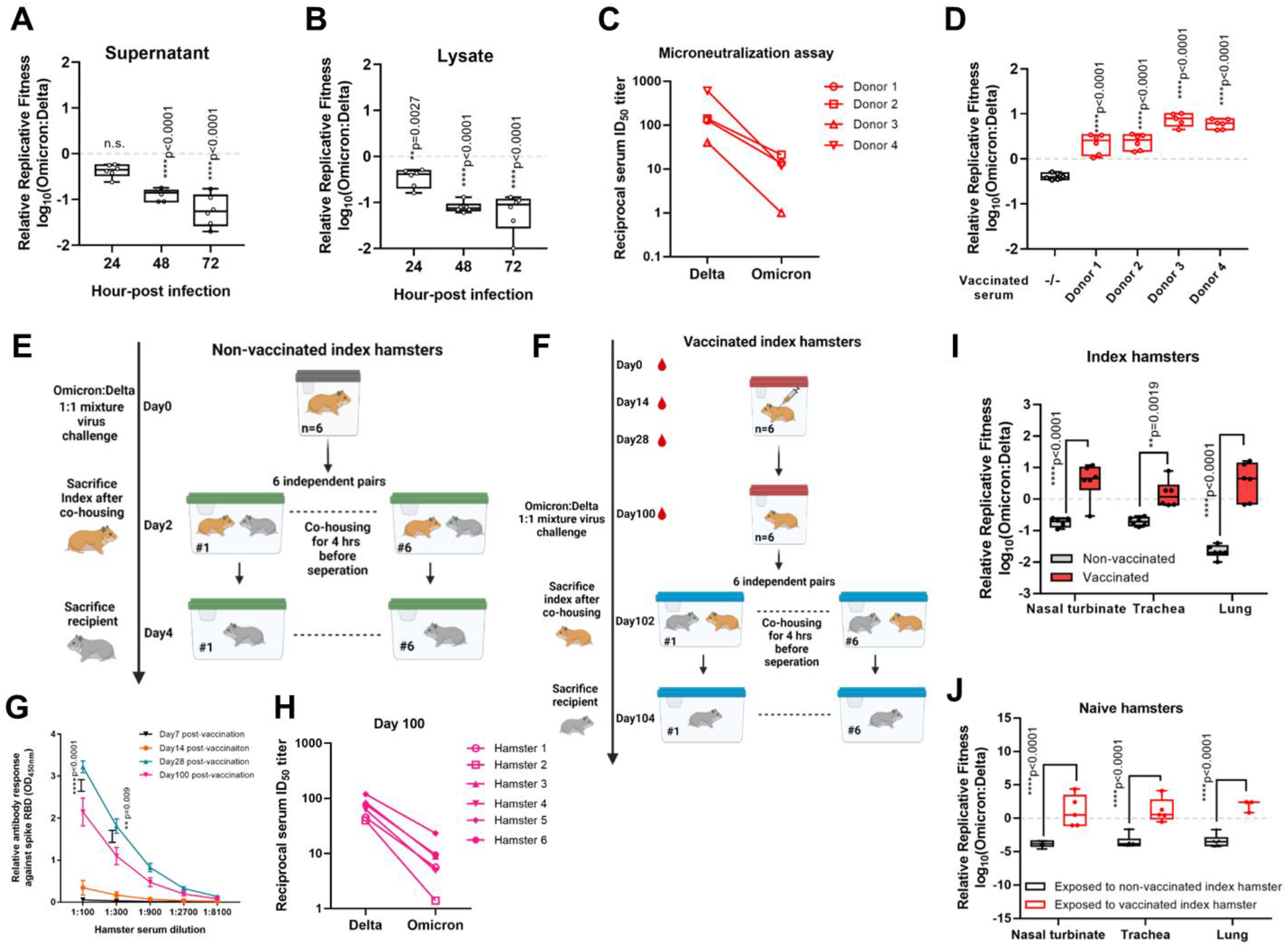
Comparative *in vitro* and *in vivo* fitness of the SARS-CoV-2 Omicron and Delta variants. **(A)** *In vitro* virus competition assay using a mixture of the Omicron and Delta variants with an initial ratio of 1:1 was inoculated onto Calu-3 cell cultures (final MOI of 0.10 for each variant). The ratios after competition in the cell culture **(A)** supernatant and **(B)** lysates were measured by qRT-PCR. **(C)** Serum samples from 4 donors at least 14 days after receiving the second dose of mRNA (Pfizer-BioNTech Comirnaty) vaccine were used to neutralize live Omicron and Delta variants, and the resulting 50% inhibitory dilution (ID_50_) were displayed. **(D)** *In vitro* virus competition with or without addition of the vaccinees’ serum samples in cell culture supernatants. Scheme of the *in vivo* competition models using **(E)** non-vaccinated and **(F)** vaccinated index hamsters. **(G)** The hamster sera (n=6) were collected at the indicated days after vaccination for detection of antibody response against spike RBD. **(H)** The neutralizing activity of the hamster serum samples collected at day 100 after vaccination against live Omicron and Delta variants. The Omicron:Delta ratio in the nasal turbinate, trachea, and lung of the **(I)** non-vaccinated and vaccinated index hamsters, and **(J)** naïve hamsters exposed to the non-vaccinated or vaccinated index hamsters. Data are mean ± standard deviations. \*\*\*\**P*<0.0001, \*\*\**P*<0.001, \*\**P*<0.01, and \**P*<0.05 by two-way ANOVA.

## DISCUSSION

Novel SARS-CoV-2 variants will continue to emerge as long as the virus maintains its wide circulation among humans and mammals. While it has recently become evident that the Omicron variant exhibits immune evasion to most existing anti-SARS-CoV-2 therapeutic monoclonal antibodies and vaccine-induced neutralizing antibodies (*9–15*), little is known about the *in vivo* pathogenicity, transmissibility, and fitness of this emerging VOC. The golden Syrian hamster model is a well-established animal model that closely mimics the clinical and virological features of COVID-19 in human, and has been widely applied for studying these aspects of SARS-CoV-2 (*19,20,22–26*).

In our pathogenicity study, we showed that while the viral load and infectious virus titer of the two variants were similar in the nasal turbinate and trachea, the Omicron variant is significantly less replicative in the lung even at the early post-infection stage (2 dpi). The infected hamsters could more efficiently restrict replication of the Omicron variant than the Delta variant by 4 dpi and 7 dpi, with consistently less cytokine/chemokine dysregulation and tissue damage from 2 dpi to 7 dpi. The lower anti-spike response in the Omicron-infected hamsters corroborates with the overall less severe disease induced by the Omicron than the Delta variant. Our *in vivo* findings corroborate with recent *in vitro* studies comparing Omicron with Delta and other SARS-CoV-2 variants. In human lung-derived Calu-3 cells, the Omicron variant shows lower replication than the Delta variant and D614G strain in in pseudovirus and/or live virus assays (*21,27*). Moreover, the Omicron spike exhibits reduced receptor binding, fusogenicity, as well as S1 subunit shedding *in vitro* (*21,27*). Interestingly, while our data indicated that the Omicron variant is generally less replicative than the Delta variant throughout the upper and lower respiratory tract, a recent preprint non-peer-reviewed study using *ex vivo* lung organ culture reported the Omicron variant showed higher replication than the Delta variant in the bronchi (*28*). This apparent difference may be caused by the different study models and conditions. It is also notable that there are significant variations in the *ex vivo* bronchi results which may have been obtained from different patient donors. Taken together, our *in vivo* findings help to explain the observations in early epidemiological studies reporting lower rates of hospitalization caused by the Omicron variant than the Delta variant. Preliminary analysis in South Africa reported that the hospitalization rate was about 30% lower among Omicron-infected patients than those infected with previous variant (*16*). Preliminary estimation in the United Kingdom also showed that the hospitalization rate of Omicron-infected patients may be 40-45% lower than that of Delta-infected patients (*29*). Importantly, we also showed that virus shedding in the oral swabs and feces were both significantly lower and shorter in the Omicron-infected than Delta-infected hamsters. This may have implications on the management and infection control of Omicron-infected patients should the same viral shedding pattern be confirmed in human.

The transmissibility of the Omicron variant is another key factor for optimizing public health control measures and predicting the evolvement of the pandemic. Recent epidemiological studies suggested that the Omicron variant may be spreading even faster (up to 4.2 times) than the Delta variant in its early stage (*30*). The estimated R_t_ of the Omicron variant in South Africa and the United Kingdom is 2.5 to 3.7, with a doubling time of approximately every 3 days (*31*). Our head-to-head comparison in this hamster model showed that that the Omicron variant has at least non-inferior, if not higher, transmissibility than the Delta variant in both contact (100% vs 100% after 4 hours of close contact) and non-contact (83.3% vs 66.7% after 6 hours of exposure) of transmission. The lack of statistical significance in our study was likely related to the use of the Delta variant which is highly transmissible itself as the comparator. It is noteworthy that the 15-20% higher rate of transmission through non-contact transmission would likely be epidemiologically significant if the same degree of enhanced transmission was found in large human populations, especially in areas with low herd immunity and relaxed public health control measures. This non-inferior or even higher transmissibility of the Omicron variant despite generally lower respiratory tract viral load than the Delta variant in our hamster model and in Calu-3 cells *in vitro* (*21*) suggests that other factors such as the efficiency of the variants to enter cells, and their ability to maintain as infectious particles in aerosols for prolonged periods should be further investigated. The functional role of the amino acid mutations found in the Omicron but not the Delta variant, for example the N501Y mutation that may be associated with enhanced transmissibility, should be further characterized (*32*).

To provide insights on whether the Omicron variant will likely replace the Delta variant to become the dominant SARS-CoV-2 strain, we compared their *in vitro* and *in vivo* fitness in cell culture and hamster models. Rather unexpectedly, the rapidly disseminating Omicron variant was consistently outcompeted by the Delta variant *in vitro* and in unvaccinated hamsters, which might be the results of its unusually high number of genetic mutations. More importantly, we showed that the Omicron variant exhibits significant fitness advantage over the Delta variant under selection pressure *in vitro* in the presence of vaccinated serum and in hamsters with waning serum neutralizing antibody level at 100 days post-vaccination with inactivated COVID-19 vaccine. Taken together, our findings suggest that the Omicron variant may outcompete the Delta variant to become the predominant SARS-CoV-2 strain especially in populations with high rates of previous infection and/or vaccination with first-generation COVID-19 vaccines eliciting ineffective neutralizing antibody response against the Omicron variant. However, our findings should be carefully interpreted and not be considered as evidence against COVID-19 vaccination. On the contrary, our findings in hamsters with waning serum neutralizing antibody at more than 3 months after vaccination are supportive of booster vaccines because recent preprint data have shown that antibody neutralization is mostly restored by mRNA vaccine booster doses (*33*).

Our study had limitations. The transmission rate of SARS-CoV-2 may vary according to different durations of exposure. In this study, we selected 6 hours of non-contact transmission to simulate the scenarios of staying with an infected index patient within the same facility for a routine business day and on medium-haul flights. It would be worthwhile to further compare the transmissibility of the Omicron and other variants after different durations of exposure in future studies. It would also be important to investigate the pathogenicity and transmissibility of the Omicron variant in additional animal models such as the hACE2-transgenic mouse and non-human primate models, as each of these animal models have their advantages and disadvantages in recapitulating human disease.

In summary, the present study shows that despite comparatively lower pathogenicity than the Delta variant, the Omicron variant undoubtedly still causes obvious clinical effects, increased viral burdens and pro-inflammatory cytokines/chemokines, as well as histopathological damages in infected hosts. Taking into consideration the Omicron variant’s higher transmissibility than the already-highly transmissible Delta variant, our findings highlight the urgent need to find next-generation COVID-19 vaccines and broad-spectrum therapeutics, as well as to tighten non-pharmaceutical measures to reduce acute and chronic disease burden (long COVID) on the public and healthcare facilities.

## MATERIALS AND METHODS

### Study design

The emerging Omicron variant has an alarmingly high number of mutations in its spike, which may not only affect its susceptibility to existing vaccines and monoclonal antibodies, but also its pathogenicity, transmission, and fitness. This study was designed to characterize these important virological attributes of the Omicron variant. To optimally place our findings into the context of the latest COVID-19 pandemic development, we performed side-by-side comparison between the Omicron variant and the currently dominant Delta variant. For the *in vivo* experiments, we adopted the well-established golden Syrian hamster COVID-19 model which closely mimics human disease (*19*). Gender- and age-matched hamsters were randomized into different experimental groups. The group sizes were chosen based on statistical power analysis and our prior experience in examining virus burdens and cytokine/chemokine profiles in hamsters. The hamsters were challenged with either SARS-CoV-2 variant and the blood and tissues were harvested for virological, cytokine/chemokine, and histopathological studies. Body weights and clinical scores were monitored until the indicated time-points. For the measurement of quantitative values, such as viral genome copies, infectious viral titers, cytokine/chemokine gene copies, body weights and clinical scores, no blinding procedures were applied to the experimentalists involved. For the histological examinations, qualified pathologists were blinded to group allocation to ensure the assessment was unbiased. All data collected was included without exclusion of outliers. All experiments in mice were individually performed in two separate occasions. For the *in vitro* competition experiment, live virus infection assays were performed to delineate the replication of the Omicron and Delta variants in human lung-derived Calu3 cells. Viral loads and infectious virus titers were quantified with RT-qPCR and plaque assays, respectively.

### Viruses and cells

SARS-CoV-2 Omicron (hCoV-19/Hong Kong/HKU-344/2021; GISAID accession number EPI_ISL_7357684) and Delta (hCoV-19/Hong Kong/HKU-210804-001/2021; GISAID: EPI_ISL_3221329) variants were isolated from respiratory tract specimens of laboratory-confirmed COVID-19 patients in Hong Kong (*9*). The inoculated cells were monitored daily for cytopathic effects by light microscopy and the cell supernatants were collected daily for qRT-PCR to assess viral load. The viruses were passaged two times before being used for the experiments. The cell lines used in this study were available in our laboratory as previously described (*34*). Calu-3 and VeroE6-TMPRSS2 cells were maintained in DMEM culture medium supplemented with 10% heat-inactivated FBS, 50 U/ml penicillin and 50 μg/ml streptomycin. All experiments involving live SARS-CoV-2 followed the approved standard operating procedures of the Biosafety Level 3 facility at The University of Hong Kong (*19,20,35*).

### *In vivo* pathogenicity study in golden Syrian hamsters

The pathogenicity of the Omicron and Delta variants were compared in the established golden Syrian hamster model for COVID-19 as we described previously (19). Briefly, male and female hamsters, aged 8-10 weeks old, were obtained from the Chinese University of Hong Kong Laboratory Animal Service Centre through the HKU Centre for Comparative Medicine Research. The hamsters were kept in biosafety level 2 housing and given access to standard pellet feed and water ad libitum. At 0 dpi, each hamster was intranasally inoculated with 100μL of DMEM containing 10^5^ PFU of SARS-CoV-2 (n=15 for each variant) under intraperitoneal ketamine (200mg/kg) and xylazine (10mg/kg) anesthesia. The body weight and clinical signs of disease of the hamsters were monitored daily. A score of 1 was given to each of the following clinical signs: lethargy, ruffled fur, hunchback posture, and rapid breathing as previously described (*36*). The hamsters were sacrificed at 2 dpi, 4 dpi, and 7 dpi (n=5 per variant per time-point) for viral load quantitation by qRT-PCR, virus titer quantitation by plaque assay, histopathological studies and immunofluorescent staining as described previously (*19,37*). Additional hamsters (n=5 per variant) were kept beyond 7 dpi for serial viral load detection in oral swabs and feces.

### SARS-CoV-2 spike RBD binding assay

Microwell plates were coated with 100 ng/well of SARS-CoV-2 RBD specific for each variant (Omicron RBD, SinoBiological #Cat:40592-V08H121 and Delta RBD, Sinobiological Cat# Cat:40592-V08H90) overnight at 4°C, followed by incubation with blocking reagent overnight at 4°C. After removal of the blocking solution, aliquots of 100 μl/well of serially diluted hamster sera from the dilution of 1:100 were added to microwell plates coated with RBD protein and incubated at 37°C for 1 h. The plates were then washed 6 times, rabbit anti-hamster horseradish peroxidase antibody (100 μL/well) at the dilution of 1:2000 was added and incubated for 30 min at 37°C. After the plates were washed 6 times, tetramethylbenzidine substrate (100 μL/well) was added. The reaction was stopped after 10 min by the addition of 0.3N sulfuric acid, and the plates were then examined in an ELISA plate at 450 nm and 620 nm.

### Contact transmission study

The contact transmission study was performed as we described previously (*19*). Briefly, Index hamsters were intranasally challenged with either the Omicron or Delta variant at 0 dpi. Twenty-four hours later, each virus-challenged index hamster was transferred to a new cage, with each cage containing one naïve hamster as a close contact. The index and naïve hamsters were co-housed for 4 hours before transferral to new cages. The index and contact hamsters were sacrificed at 2 dpi and 2 days post-exposure, respectively. Hamsters with cycle threshold (Ct) value <40 in either nasal turbinate or lung were considered as infected. The experiment was repeated twice, each time with 6 index (n=3 for Omicron and n=3 for Delta each time) and 6 naïve hamsters (n=3 for Omicron and n=3 for Delta each time).

### Non-contact transmission study

The transmissibility of the Omicron and Delta variants among hamsters through non-contact transmission was compared in our established closed system with unidirectional airflow in two independent experiments (*20*). In each experiment, 6 index hamsters (n=3 for each variant), each housed in a separate cage (Marukan Co., Ltd., Osaka, Japan) inside isolators (Tecniplast SpA, Varese, Italy), were intranasally challenged with 100μL of Dulbecco’s Modified Eagle Medium (DMEM) containing 10^5^ plaque-forming units of either variant under intraperitoneal ketamine and xylazine anaesthesia at 0 dpi. After 24h, 36 naïve hamsters were transferred to cages adjacent to the cages housing the index hamsters (1:6 ratio). The index and exposed naïve hamsters were removed from the non-contact transmission system and transferred to separate new cages 6 hours later. The hamsters were then sacrificed at 2 dpi (index) or 2 days post-exposure (naïve) for organ tissue collection. Hamsters with Ct value <40 in either nasal turbinate or lung were considered as infected. The experiment was repeated twice, with a total of 12 index and 72 naïve hamsters.

### *In vitro* virus competition assay

Approximately 3×10^5^ cells were seeded onto each well of 12-well plates and cultured at 37°C, 5% CO_2_ for 16 h. Equal PFUs of two variants (1:1 ratio) were inoculated onto Calu-3 cells at a final MOI of 0.10 for each variant. The mixed viruses were incubated with the cells at 37°C for 2h. After infection, the cells were washed twice with PBS to remove residual viruses. One milliliter of culture medium was added into each well. At each time-point, 350μl cell culture lysate and/or supernatant was collected for RNA extraction. Ratios of Omicron:Delta RNA were determined via RT-PCR with quantification of Sanger peak heights or via qRT-PCR using specific primer and probes. All samples were stored at −80°C until analysis.

### *In vivo* virus competition models

To evaluate the competitive fitness of the Omicron and Delta variants *in vivo*, virus competition experiments were performed using non-vaccinated and vaccinated hamsters. In the non-vaccinated hamster model, six index hamsters were infected each intranasally with 10^5^ PFU of a mixture of both viruses (1:1 ratio). At 2 dpi, each index hamster was co-housed with one naïve hamsters for 4 hours in six independent lines of naïve animals. Total RNA was extracted from the nasal turbinate, trachea, and lung tissues, of the hamsters at 2 dpi or 2 days post-exposure (exposed naïve) using RNeasy kit (Qiagen) for viral load quantification and calculation of the Omicron:Delta ratio. In the vaccinated hamster model, each index hamster was intramuscularly administered with 3μg per dose of inactivated vaccine at day 0 and boosted at day 14, respectively. To generate the inactivated virus vaccine, virus-containing supernatant [SARS-CoV-2 virus HKU-001a strain, GenBank accession number: MT230904)] was harvested and inactivated using 0.2% formaldehyde solution in PBS for 5 days. Cell cytopathic effects (CPE) were monitored to validate complete inactivation using Vero cells. Hamster sera were collected at day 0, day 14, day 28, and day 100 for neutralization antibody titration using enzyme-linked immunosorbent assay and micro-neutralization assay. At day 100 post-vaccination, the vaccinated index hamsters were infected (ie: 0dpi) and the competition experiments were performed as described for the non-vaccinated hamster model.

### Validation of competition assay

The experiments were performed as previously described (*26*). To validate the consistency and accuracy of the competition assay in both system, the Omicron and Delta variants were mixed at ratios of 10:1, 5:1, 3:1, 1:1, 1:3, 1:5, and 1:10 based on their PFU titres (total 10^5^ PFU viruses), or mixed with 10^6^, 10^5^, 10^4^, 10^3^, and 10^2^ PFU of the two variants at a ratio of 1:1. The total RNA of these mixed variants was isolated and amplified by RT-PCR followed by Sanger sequencing, or directly analysed by one-step RT-qPCR using strain-specific primers and probes as mentioned below. The Omicron to Delta ratio was calculated by the peak heights of Sanger sequencing, i.e. Spike_R493 for Omicron (CGA) and Spike_Q493 for Delta (CAA), or the copy numbers by the RT-qPCR system targeting a spike region of Omicron and Delta. Data were analysed by linear regression with correlation coefficients (r) and significance (P).

### Quantification of RNA ratios between Omicron and Delta variants

To measure the ratios Omicron to delta for competition assays, both qRT-PCR and Sanger sequencing were performed. Real-time one-step qRT-PCR was used for quantitation of Omicron and Delta spike gene copy within the same input template after RNA extraction. The QuantiNova Probe RT-PCR kit (Qiagen) was used with a LightCycler 480 Real-Time PCR System (Roche) as previously described (*38*). Each 20μL reaction mixture contained 10μL of 2×QuantiNova Probe RT-PCR Master Mix, 0.8μL of RNase-free water, 0.2μL of QuantiNova Probe RT-Mix, 0.8μL each of 20μM forward and reverse primers, 0.4μL each of 10μM probes, and 5μL of extracted RNA as the template. Reactions were incubated at 45°C for 10 min for reverse transcription, 95°C for 5 min for denaturation, followed by 45 cycles of 95°C for 5s and 55°C for 30s. Signal detection and measurement were taken in each cycle after the annealing step. The cycling profile ended with a cooling step at 40°C for 30s. Primers and probes to be used are as below: Omi-S_F: ACAAACCTTGTAATGGTGTTGC; Omi-S_R: TACTACTACTCTGTATGGTTGGTG; Omi-S_probe: Cy5-CGATCATATAGTTTCCGACCCACTT–IAbRQSp; Delta-S_F: GCAAACCTTGTAATGGTGTTGA; Delta-S_R: GTACTACTACTCTGTATGGTTGGTA; Delta-S_probe: FAM-CAATCATATGGTTTCCAACCCACTA –IABkFQ. For Sanger sequencing, RT-PCR was conducted using a SuperScript™ III One-Step RT-PCR kit (Invitrogen, Carlsbad, CA, USA) to amplify the extracted viral RNA (Qiagen). Relative replicative fitness values for omicron virus compared to delta virus were analysed according to *w=*(*f0/i0*), where *i0* was the initial Omicron/Delta ratio and *f0* was the final Omicron/Delta ratio after competition. Sanger sequencing (initial time-point T0) counts for each variant being compared were based upon average counts over repeated samples of inoculum per experiment, and post-infection (time-point T1) counts were taken from samples of individual subjects. To model *f0/i0*, the ratio T0/T1 was determined for each subject in distinct strain groups.

### Statistical analysis

All data were analysed with GraphPad Prism software (GraphPad Software, Inc). Body weight losses were compared using two-way ANOVA. Two-way ANOVA or Student’s t-test was used to determine significant differences in viral loads and titers. P<0.05 was considered statistically significant.

### Ethical approvals

The animal experiments were approved by the Institutional Review Board of The University of Hong Kong Committee on the Use of Live Animals in Teaching and Research (CULATR) and the use of clinical specimens was approved by the Institutional Review Board of the University of Hong Kong / Hospital Authority Hong Kong West Cluster.

### Illustrations

The hamster illustrations and schematic figures were created with BioRender software (https://biorender.com/).

## Supplementary Materials

## Author contributions

Conceptualization: S.Y., K.-Y.Y., J.F.-W.C.

Methodology: S.Y., D.-Y.J., J.F.-W.C.

Investigation: S.Y., Z.W.Y., R.L., K.T., A.Z., G.L., C.P.O., V.K.-M.P., C.C.-S.C., B.W.-Y.M., Z.Q., Y.X., H.S., J.O.-L.T., T.T.-T.Y., K.K.-H.C., C.C.Y.C., J.-P.C., C.L., C.C.-Y.Y., L.L., K.K.-W.T., H. Chen, H. Chu, J.F.-W.C.

Visualization: S.Y., J.F.-W.C.

Funding acquisition: G.L., H. Chen, D.-Y.J., K.-Y.Y., J.F.-W.C.

Project administration: S.Y., D.-Y.J., K.-Y.Y., J.F.-W.C

Supervision: S.Y., D.-Y.J., K.-Y.Y., J.F.-W.C

Writing – original draft: S.Y., J.F.-W.C

Writing – review & editing: S.Y., D.-Y.J., K.-Y.Y., J.F.-W.C

## Data availability

Complete sequences of the SARS-CoV-2 Omicron (hCoV-19/Hong Kong/HKU-344/2021; GISAID accession number EPI_ISL_7357684) and Delta (hCoV-19/Hong Kong/HKU-210804-001/2021; GISAID: EPI_ISL_3221329) variants available through GISAID. Other supporting raw data are available from the corresponding author upon reasonable request. Source data are provided with this paper.

---

**Fig. S1.**
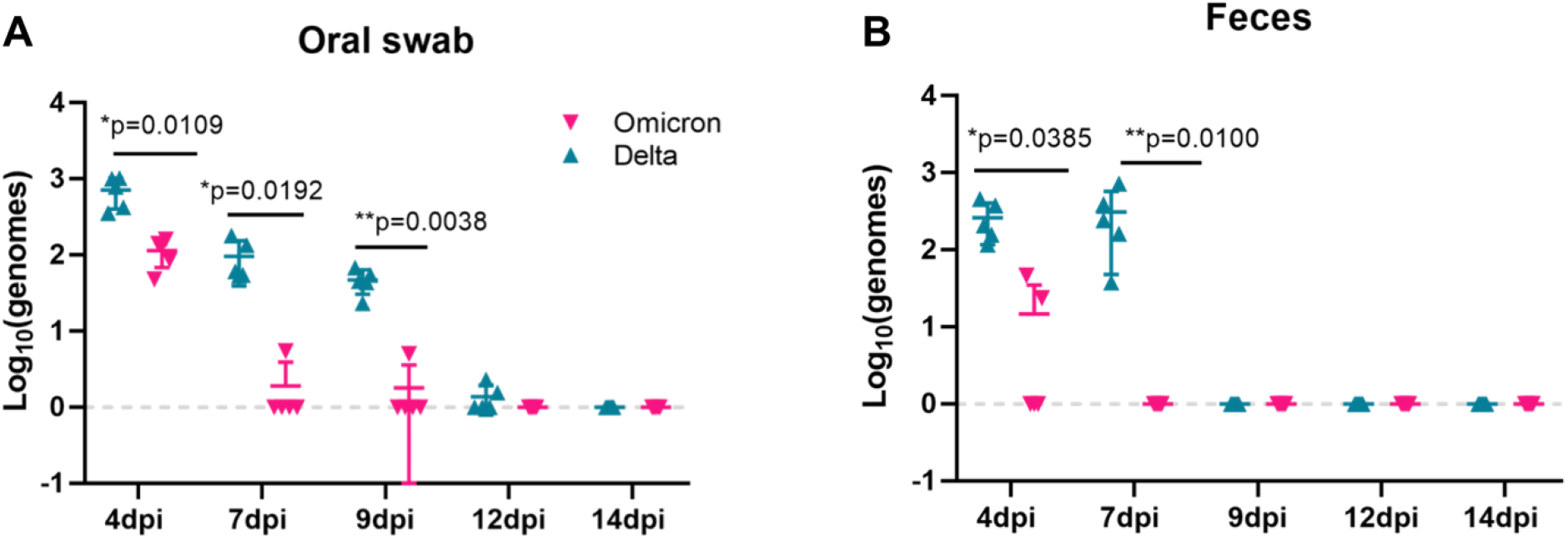
Virus shedding in the oral swabs and feces of the Omicron-infected and Delta-infected hamsters. **(A)** Oral swabs and **(B)** feces were serially collected from the Omicron-infected and Delta-infected hamsters at the indicated time-points for viral load detection by quantitative reverse transcription-polymerase chain reaction. The dotted line represented the limit of detection. Data are mean ± standard deviations. n = 5 biological replicates per variant. \*\**P*<0.01 and \**P*<0.05 by two-way ANOVA.

**Fig. S2.**
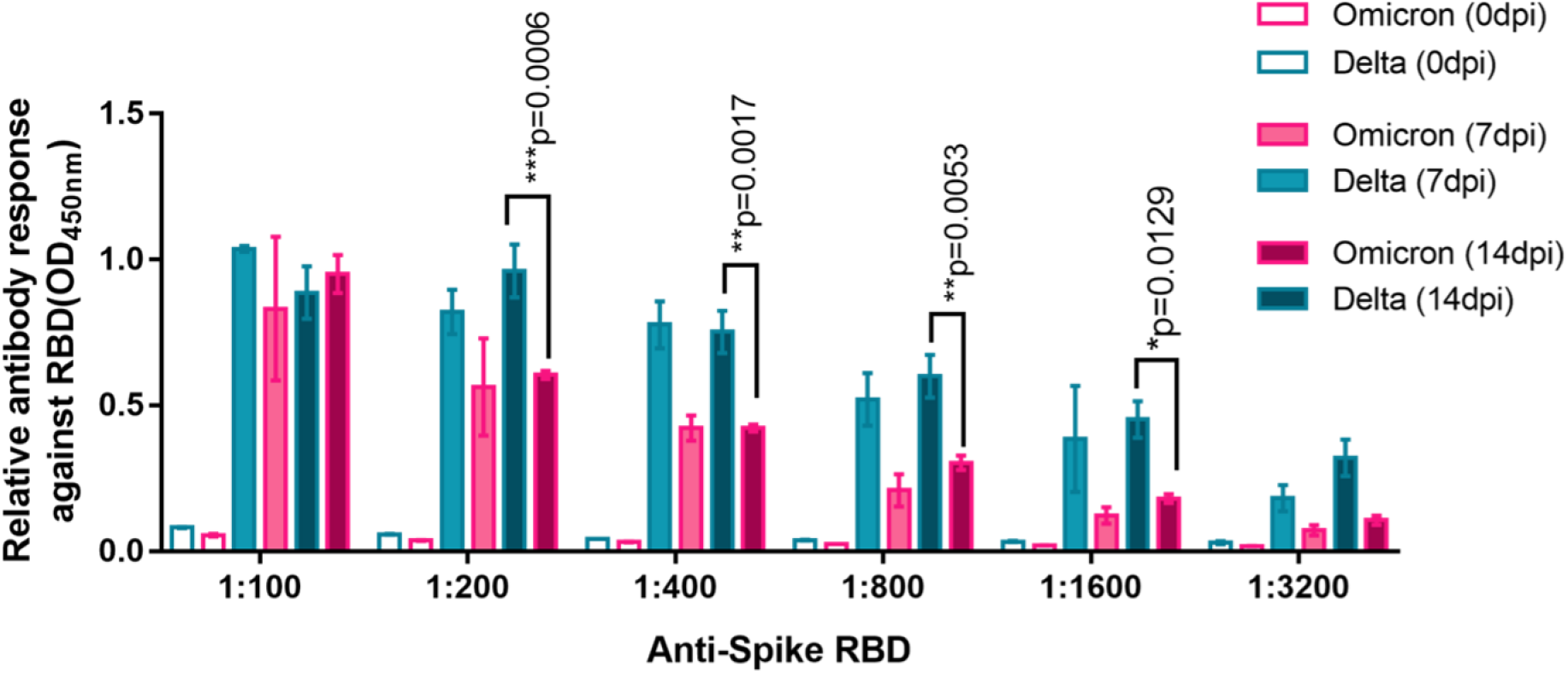
Variant-specific anti-spike receptor-binding domain antibody response of the Omicron-infected and Delta-infected hamsters. Microwell plates were coated with 100 ng/well of SARS-CoV-2 RBD specific for each variant (Omicron RBD, SinoBiological #Cat:40592-V08H121 and Delta RBD, Sinobiological Cat# Cat:40592-V08H90) overnight at 4°C, followed by incubation with blocking reagent overnight at 4°C. After removal of the blocking solution, aliquots of 100 μl/well of serially diluted hamster sera from the dilution of 1:100 were added to microwell plates coated with RBD protein and incubated at 37°C for 1 h. The plates were then washed 6 times, rabbit anti-hamster horseradish peroxidase antibody (100 μL/well) at the dilution of 1:2000 was added and incubated for 30 min at 37°C. After the plates were washed 6 times, tetramethylbenzidine substrate (100 μL/well) was added. The reaction was stopped after 10 min by the addition of 0.3N sulfuric acid, and the plates were then examined in an ELISA plate at 450 nm and 620 nm. n = 5 biological replicates per variant. \*\*\*\**P*<0.0001, \*\*\**P*<0.001, \*\**P*<0.01, and \**P*<0.05 by two-way ANOVA.

**Fig. S3.**
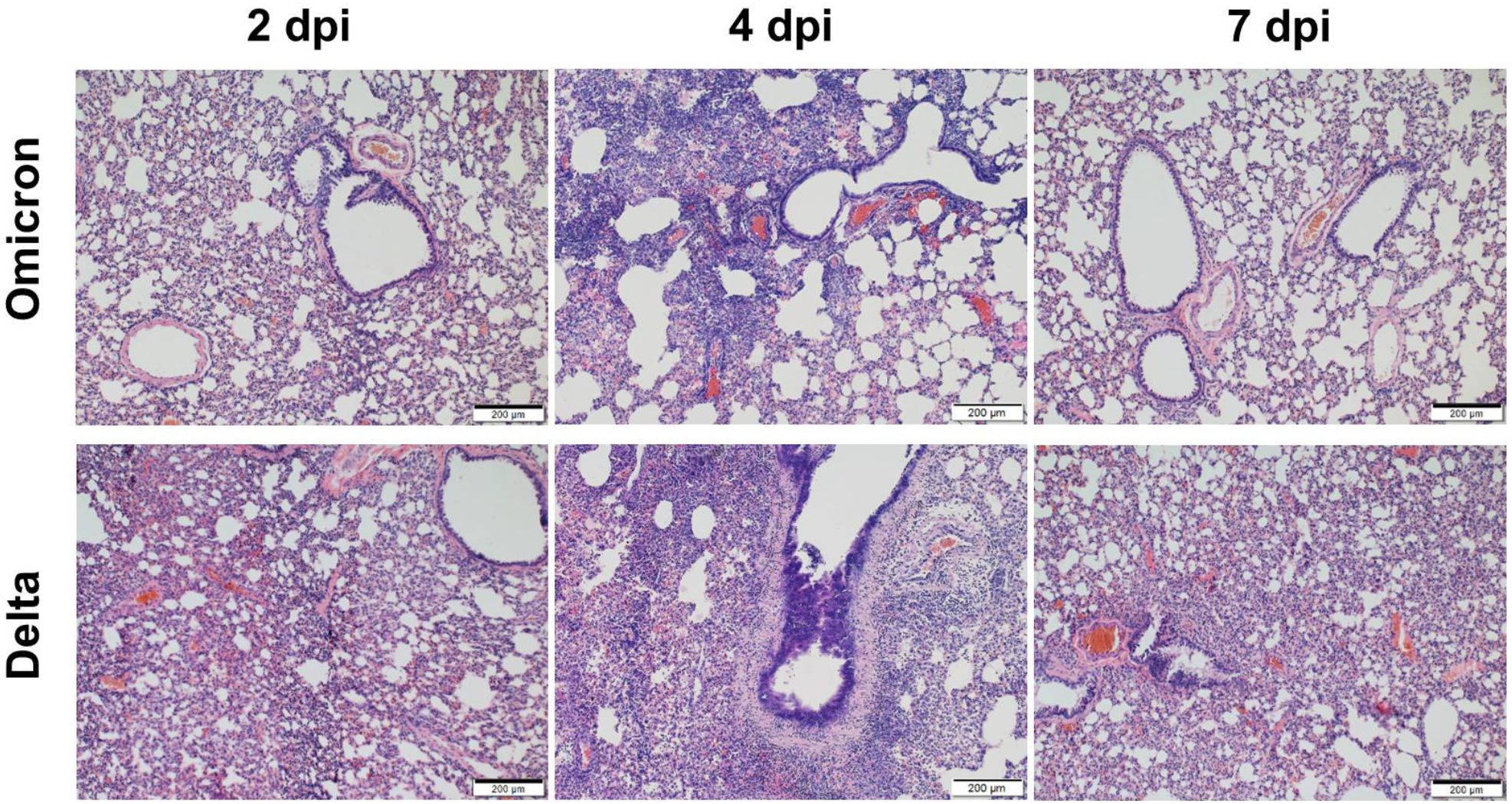
Representative histopathological findings of the lung of Omicron-infected and Delta-infected hamsters. Representative haematoxylin and eosin-stained lung sections of the Omicron-infected and Delta-infected hamsters were collected at 2 days post-infection (dpi), 4 dpi, and 7 dpi. At 2 dpi and 4 dpi, the Delta-infected hamsters showed more diffuse peribronchiolar, perivascular, and alveolar inflammatory infiltrates, bronchiolar epithelial destruction, and alveolar congestion than the Omicron-infected hamsters. At 7 dpi, the histopathological changes have mostly resolved in the lungs of the Omicron-infected hamsters, while the Delta-infected hamsters still demonstrated blood vessel congestion and alveolar wall inflammatory infiltration. Scale bars = 200μM.

**Fig. S4.**
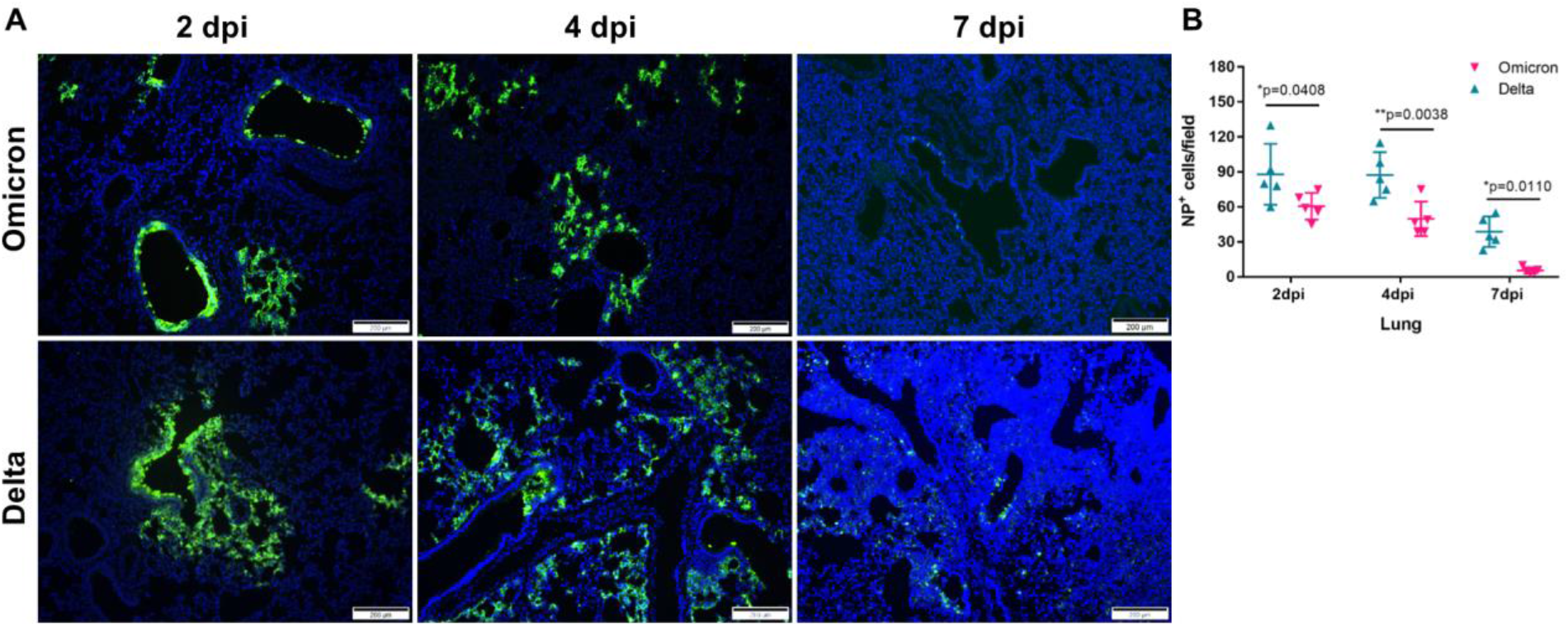
Representative immunofluorescence staining findings of the lung of Omicron-infected and Delta-infected hamsters. **(A)** Immunofluorescence staining of the lung sections of Omicron-infected and Delta-infected hamsters at 2 days-post infection (dpi), 4 dpi, and 7 dpi. SARS-CoV-2 nucleocapsid (NP, green) and cell nuclei (blue) were stained. These representative images were selected from a pool of 15 images captured in 5 hamsters per variant. **(B)** NP-positive cells per 50× field per lung section of each hamster. Scale bars = 200 μm. Data are mean ± standard deviations. n = 5 biological replicates per variant. \*\**P*<0.01 and \**P*<0.05 by two-way ANOVA.

**Fig. S5.**
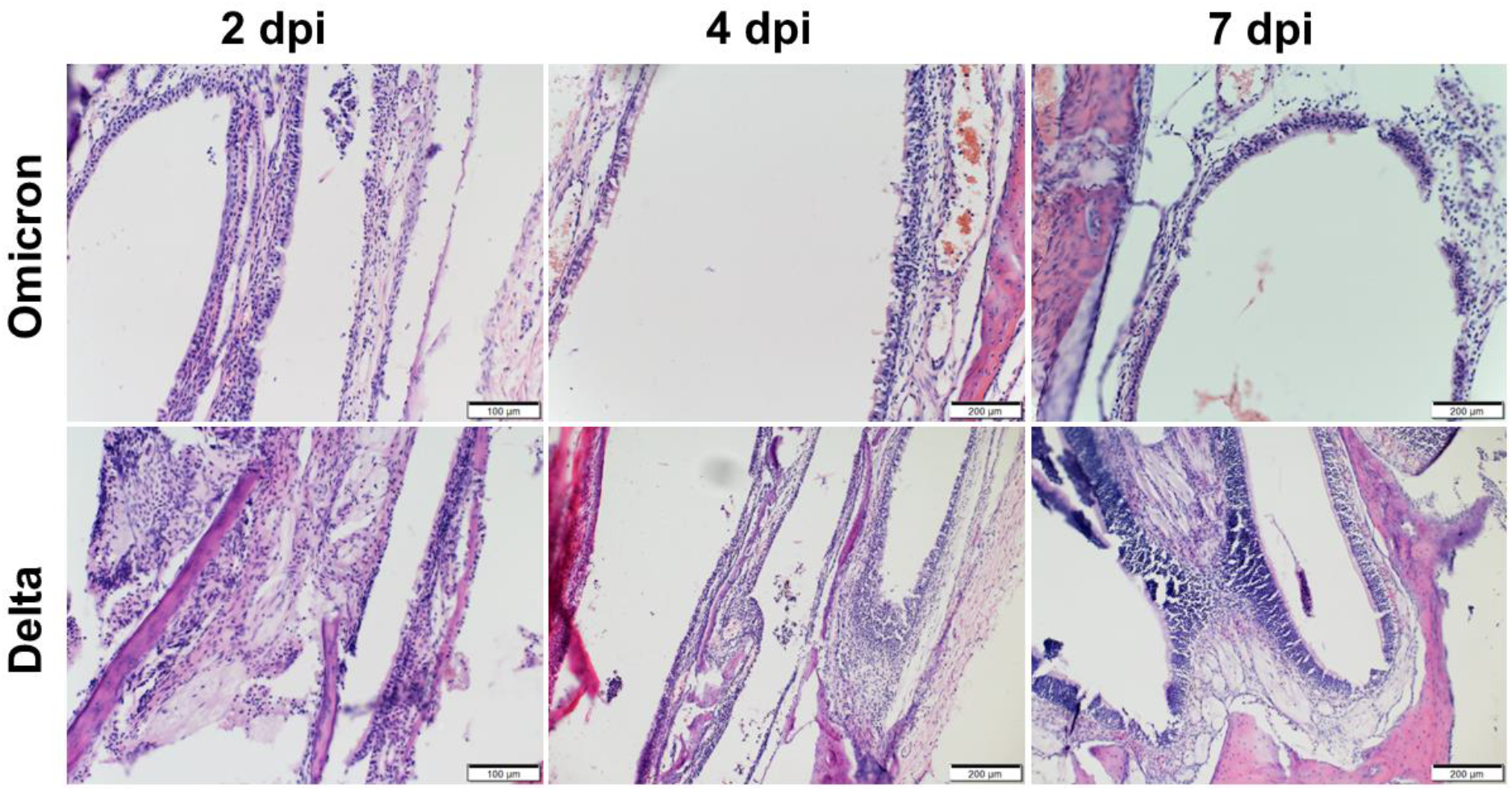
Representative histopathological findings of the nasal turbinate of Omicron-infected and Delta-infected hamsters. Representative haematoxylin and eosin-stained lung sections of the Omicron-infected and Delta-infected hamsters were collected at 2 days post-infection (dpi), 4 dpi, and 7 dpi. At 2 dpi, submucosal infiltration and destruction of the epithelium were observed in both groups of hamsters. At 4 dpi, the Delta-infected hamsters showed severe destructive changes of the respiratory and olfactory epithelia, while the Omicron-infected hamsters showed intact respiratory epithelium with mild intra-epithelial inflammatory infiltrates. At 7 dpi, the histopathological changes have mostly resolved in the Omicron-infected hamsters, while the Delta-infected hamsters still demonstrated some residual inflammation. Scale bars = 100μM or 200μM.

**Fig. S6.**
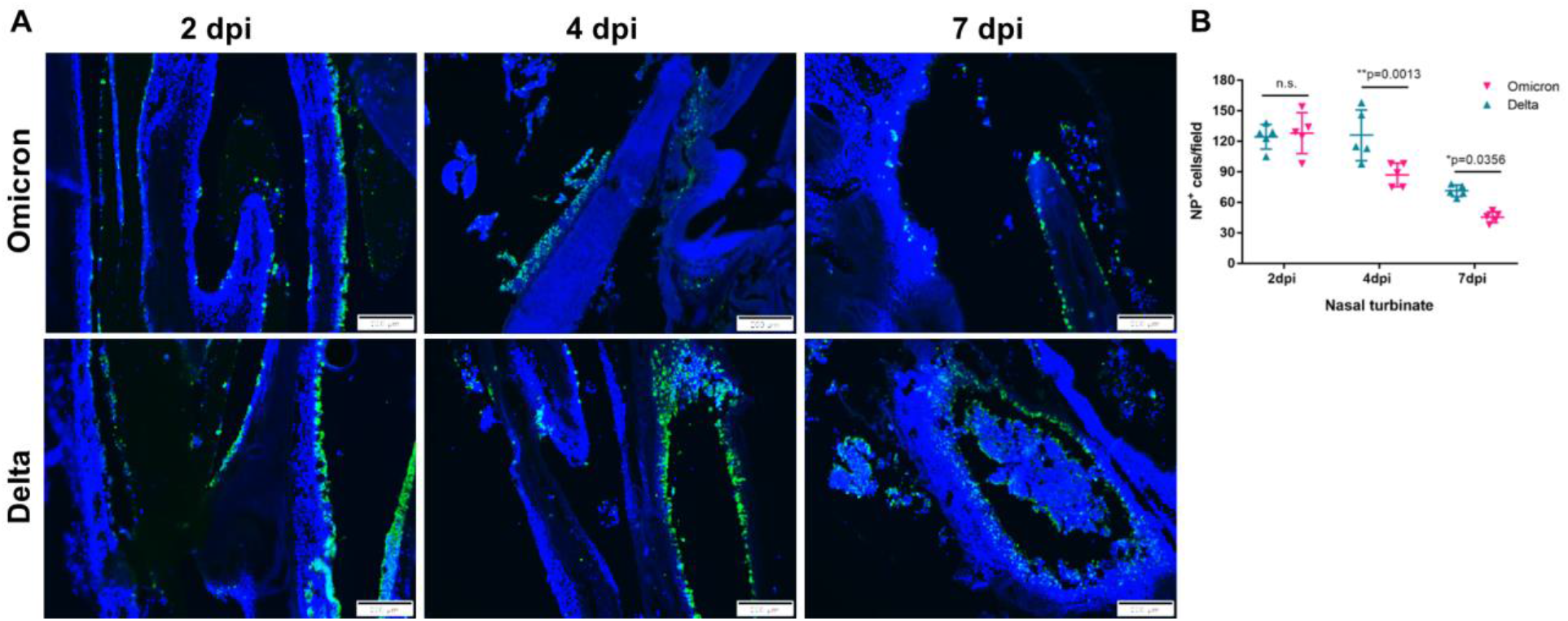
Representative immunofluorescence staining findings of the nasal turbinate of Omicron-infected and Delta-infected hamsters. **(A)** Immunofluorescence staining of the nasal turbinate sections of Omicron-infected and Delta-infected hamsters at 2 days-post infection (dpi), 4 dpi, and 7 dpi. SARS-CoV-2 nucleocapsid (NP, green) and cell nuclei (blue) were stained. These representative images were selected from a pool of 15 images captured in 5 hamsters per variant. **(B)** NP-positive cells per 50× field per nasal turbinate section of each hamster. Scale bars = 200 μm. Data are mean ± standard deviations. n = 5 biological replicates per variant. \*\**P*<0.01 and \**P*<0.05 by two-way ANOVA.

**Fig. S7.**
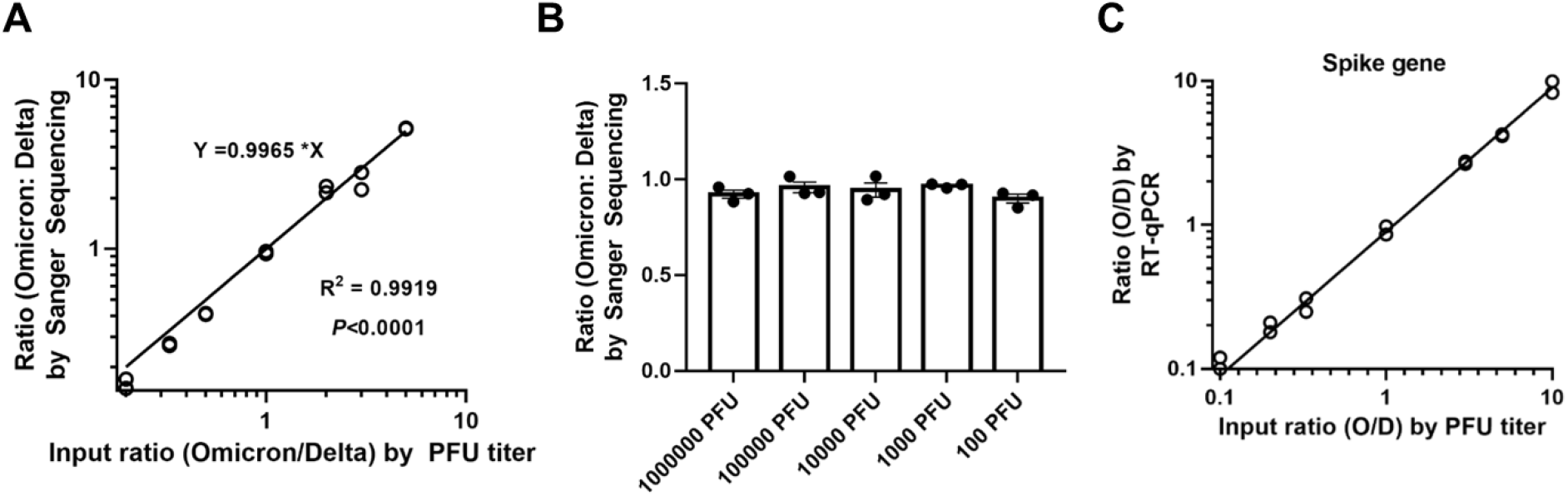
Validation of the competition assay. **(A)** The correlation between input PFU ratios and output RT-PCR amplicon ratios determined by Sanger sequencing, i.e. R493 for Omicron (CGA) and Q493 for Delta (CAA). The Omicron and Delta variants were mixed at PFU ratios of 10:1, 5:1, 3:1, 1:1, 1:3, 1:5, or 1:10. Total RNA of the mixture was extracted and amplified by RT-PCR. The R493/Q493 ratio was calculated by the peak heights of Sanger sequencing. Data were analyzed by linear regression with correlation coefficients (r) and significance (p). The symbols represent individual replicates and the solid line represents the fitted line. Data were derived from an experiment conducted in duplicate. **(B)** Assay range evaluation. The ratio of Omicron:Delta variant mixture determined by Sanger sequencing was consistent in a wide range of virus amounts. The two variants were mixed at 1:1 ratio. The total titers of the mixed virus stock were 10^2^, 10^3^, 10^4^, 10^5^, and 10^6^ PFU. The total RNA of the virus mixture was isolated and amplified by RT-PCR. The R493/Q493 ratio was calculated by the peak heights from Sanger sequencing. The symbols represent individual replicates, bar heights represent the means, and error bars represent standard deviations. Data are derived from an experiment conducted in triplicate. **(C)** Validation of the competition assay by spike-targeting qRT-PCR that differentiates the Omicron and Delta variants.

